# LEMMI: A continuous benchmarking platform for metagenomics classifiers

**DOI:** 10.1101/507731

**Authors:** Mathieu Seppey, Mosè Manni, Evgeny M. Zdobnov

## Abstract

Studies of microbiomes are booming, as well as the diversity of computational tools to make sense out of the sequencing data and the volumes of accumulated microbial genotypes. LEMMI (https://lemmi.ezlab.org) is a novel concept of a benchmarking platform of computational tools for metagenome composition assessments that introduces: a continuous integration of tools, their multi-objective ranking, and an effective distribution through software containers. Here, we detail the workflow and discuss the evaluation of some recently released methods. We see this platform eventually as a community-driven effort: where method developers can showcase novel approaches and get unbiased benchmarks for publications, while users can make informed choices and obtain standardized and easy-to-use tools.

## Introduction

Metagenomics has made possible the study of previously undiscovered uncultured microorganisms. To probe the vast hidden microbial landscape, we need effective bioinformatics tools, notably taxonomic classifiers for binning the sequencing reads (i.e. grouping and labeling) and profiling the corresponding microbial community (i.e. defining the relative abundance of taxa). This is computationally challenging, requiring time and resources to query shotgun sequencing data against rapidly expanding genomic databases. Users have to choose a solution among the plethora of methods that are being developed in a quest for accuracy and efficiency, for instance lowering the runtime and memory usage by reducing the reference material while maintaining the representative diversity^1,2^. To date, at least one hundred published methods can be identified^3^, and new developments that may revolutionize the field cohabit with re-implementations of already explored strategies, complicating methods selection, fragmenting the community of users, and hindering experimental reproducibility. Papers describing these methods fall in the “self-assessment trap”^4^, in which all published methods are the best on carefully selected data, giving no indication to potential users about general performances, which should be a prerequisite before considering any case-specific improvements or novel features. Independent comparative benchmarking studies^5–9^ and challenges^10^ are a major step towards a fair assessment of methods. They can also promote developments dedicated to specific technologies or problems^11^ and have brought valuable benchmarking resources^12–14^. While results in such publications are extensively discussed by the authors upon release, their content cannot be reinterpreted according to specific individual needs. More importantly, tools that were overlooked or made available after the list of methods was finalized cannot be added, maintaining a permanent uncertainty about the true state of the art and delaying the benefits that innovative developments bring in a fast-evolving field. To overcome these limitations, there is a need for a different workflow that enables continuous and generalizable comparisons of individual methods or multi-step pipelines while considering multiple objectives and their computational costs (e.g. accuracy versus memory usage).

Recommendations for efficient “omics” tool benchmarking^15–17^ include using containers (i.e. isolated software packages) to allow reproducibility and long-term availability of tools^18^, reporting computational resources consumption (available hardware limits the choice to otherwise less efficient methods), exploring parameters, and avoiding rankings based on a unique metric. In the particular case of taxonomic classification, a benchmark that uses the same reference for all methods is necessary to perform a valid evaluation of their respective algorithms, in addition to evaluating the variety of available databases. In line with this, we introduce LEMMI, standing for “A <u>L</u>ive <u>E</u>valuation of Computational <u>M</u>ethods for <u>M</u>etagenome Investigation” (Figure 1), a web-based platform that hosts in its infrastructure, currently, a semi-automated benchmarking pipeline whose first available component presented here focuses on assessing taxonomic profilers and binners. Its novel workflow explicitly addresses key problems for the community: (i) closing the time gap between benchmarking publications by continuously evaluating new methods, (ii) allowing heterogeneous tools and pipelines to be evaluated together with a multi-objective exploration and ranking of their performances, (iii) reporting computational resources, (iv) exploring parameters and references, (v) facilitating the dissemination of easy-to-use software packages, and (vi) producing evaluations in a neutral and controlled environment to ensure published methods have a reliable benchmark on generic problems and unified hardware. Our solution can complement and support analyses of specific cases usually discussed in self-evaluations or discussed periodically in comparative studies. We describe below the advantages of the LEMMI workflow and discuss novel results from which conclusions about several methods released recently can be drawn.

**Fig. 1.**
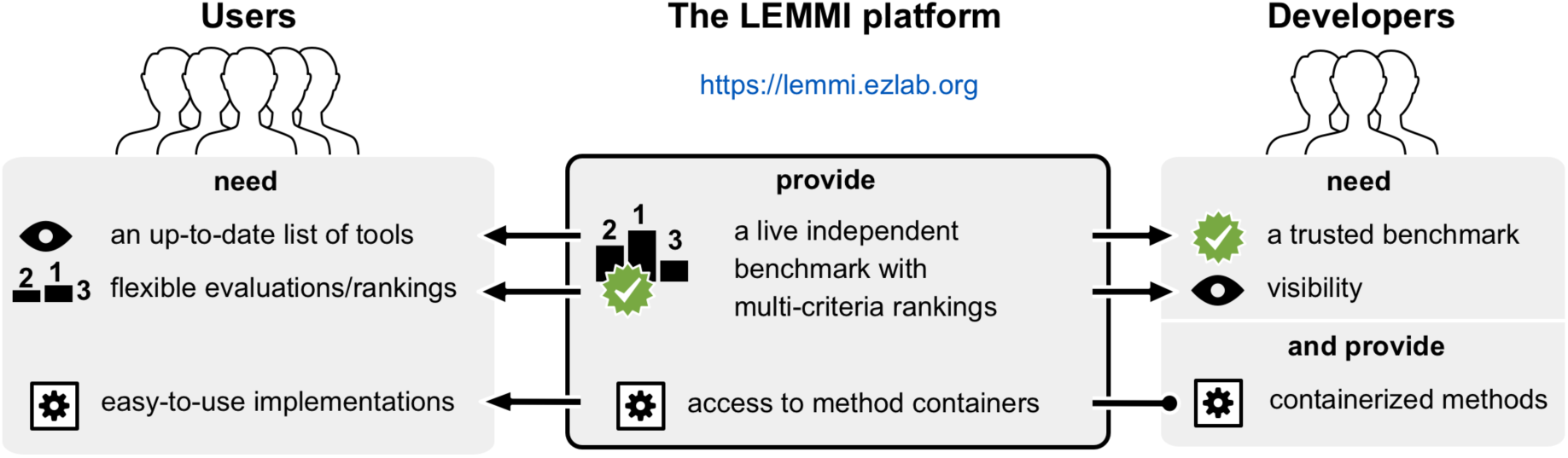
Connecting users and developers through benchmarking. The LEMMI platform facilitates the access to up-to-date implementations of methods for taxonomic binning and profiling. LEMMI creates a link between developers and users as it provides independent and unbiased benchmarks that are valuable to both. Developers need comparative evaluations to keep improving the methodology and to get their work published in peer-reviewed journals. Method users need a resource that keeps track of new developments and provides a flexible assessment of their performances to match their experimental goals, resources, and expectations. The containerized approach of LEMMI guarantees the transfer of a stable and usable implementation of each method from the developer to the final user, as they appear in the evaluation.

## Results

### A fully containerized evaluation

LEMMI inputs consist of methods wrapped in containers resembling a previously suggested format, bioboxes^14^, that can be prepared by the LEMMI development teams, and envisioned to be contributed by method developers themselves. In the latter case, containers can be exchanged through public channels (e.g. https://hub.docker.com) to be loaded and run on the platform. In addition, evaluated containers can be downloaded by users from the LEMMI website along with their building sources, unless the method is commercialized and/or not under open source licensing. LEMMI is the first solution in its field to compel containerization to manage a benchmarking workflow (Supplementary Figure 1a) and therefore favor a simple access to evaluated tools. The benefits of such an approach regarding systematic re-evaluation (Supplementary Figure 1b) and re-usability are likely greater in the long term than the flexibility offered by mere results submission. As solutions that facilitate automated conversion between distribution channels are being explored (for instance from BioConda to biocontainers^19^), establishing standardized benchmarking processes that rely exclusively on these channels to obtain all methods ensure that the otherwise subjective question of installability is addressed unambiguously.

### A dynamic interface to explore alternative objectives

Making informed decisions when designing analyses or pipelines requires a thorough exploration of the parameter space of available algorithms. Multiple runs of a container enable such exploration within the platform. LEMMI does not segregate methods into profilers and binners by conducting separate evaluations, as many tools now provide both features or have extra scripts that blur the demarcation (e.g. kraken^20,21^ is a read classifier that can be associated with its companion tool bracken^22^ to produce a valid taxonomic abundance profile). Instead, LEMMI is “results-oriented”, so comparisons can be made between multi-functional tools or their combinations to generate profiles, bins, or both, in a single run that informs on the full potential of a method. These two-dimensional results can be visualized from different perspectives through a dynamic multi-criteria ranking where users can put the emphasis on the metric they recognize as important (Figure 2-3, Supplementary Figure 2-3, 5-9). Tradeoffs can be found to maximize the efficiency of a single tool or pipeline for both tasks.

**Fig. 2.**
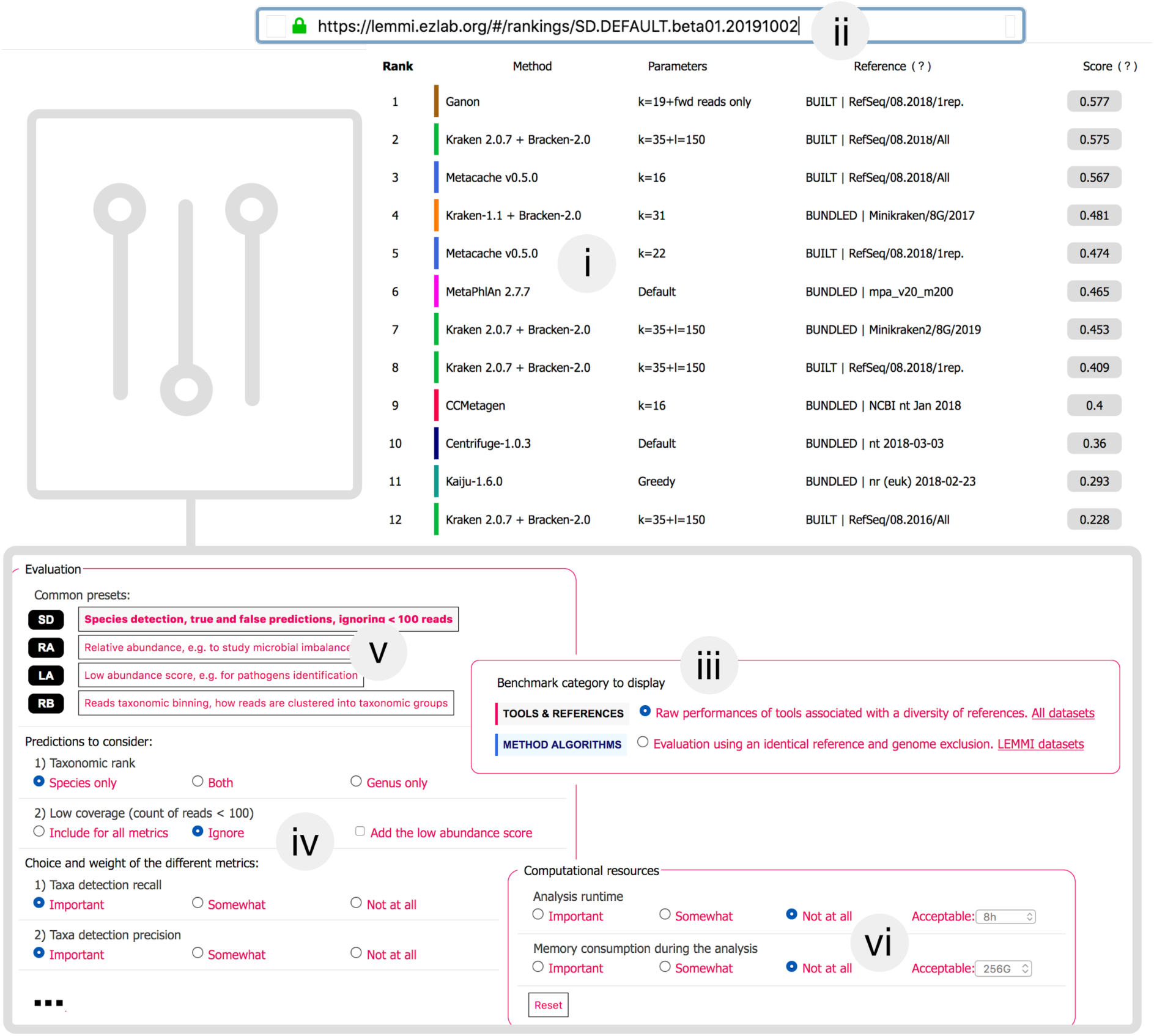
LEMMI web interface. (i) The LEMMI users obtain a list of entries suited to their needs through the dynamic ranking interface that allows them to select and weight the criteria that are important. The ranking visible here shows the performances when identifying species while ignoring taxa under 100 reads, balancing precision and recall. The score assigned to each entry visible in LEMMI rankings averages selected metrics over all tested datasets. (ii) A unique “fingerprint” allows custom rankings to be shared and restored at any time on the LEMMI platform through the address bar. (iii) The benchmark category can be selected in the dashboard. (iv) Prediction accuracy metrics can be chosen along with their importance (Weight for “Important” is 3, “Somewhat” is 1, and “Not at all” is 0). This will cause the list to be updated with the corresponding scores. (v) Several presets corresponding to common expectations are available. (vi) Computational resources and the time required to complete the analysis can be included as an additional factor to rank the methods.

**Fig. 3.**
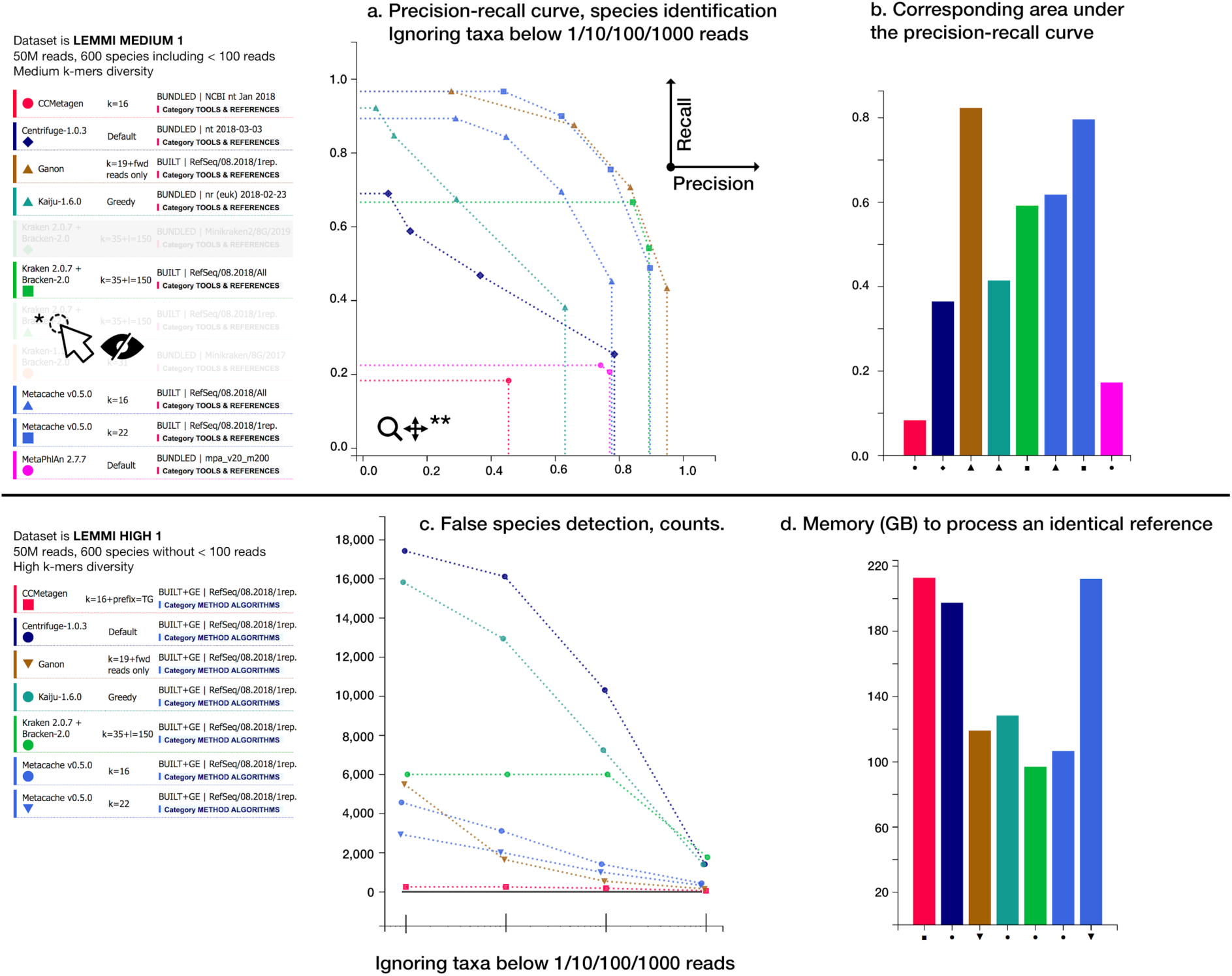
“Details by datasets” interface. Plots for up to 17 metrics are available for each dataset and the LEMMI user can toggle each line representing a method associated with a reference and specific parameters individually (*). Plots can be zoomed in to disentangle overlapping points and permit focused interpretations (**). The pages exist for all taxonomic ranks investigated, i.e. genus and species in the release beta01. The illustration shows in the upper panel (a) the precision-recall curve in species identification of a mix of methods using references freely provided or built using the current maximal capacity of LEMMI of 245GB, and (b) the corresponding area under the precision-recall curve. The lower panels show results when assessing methods using an identical built reference, with panel (c) illustrating how LEMMI assesses the precision improvement by filtering low abundance taxa, and (d) reporting the peak of memory necessary to construct the reference.

### Pre-packaged and built reference databases

Some method implementations provide pre-packaged reference databases while others provide scripts to generate them. LEMMI evaluates these pre-packaged references to include methods whose value lies in providing a curated database (e.g. marker genes^23^) and to keep track of reference genome catalogs available to method users. However, as the use of different reference databases is likely a major source of result discrepancies, methods accepting any nucleotide or protein files to construct their database can integrate the corresponding scripts in the container to be used as part of the benchmark. This enables LEMMI to report the resources (i.e. memory, runtime) required to build a reference in addition to those of the analysis, and assess the algorithm underlying the method independently from the taxa that constitute its default reference database. To provide a continuous evaluation that is unaffected by the publication of new genomic sequences, LEMMI maintains an in-house repository currently based on all bacterial and archaeal RefSeq^24^ assemblies. However, only entries having both nucleotide and corresponding protein files are kept to offer the two types of molecules as a possible reference with equal representation (hereafter, the LEMMI/RefSeq repository). It can be subsampled by publication date, assembly states, or a fixed number of representatives per taxonomic identifier (taxid). As for parameters, this enables for each method the exploration of references, for instance by restraining the source material to what was available at a given date (Supplementary Figure 3) or to specific criteria (e.g. assembly states being only “Complete Genome”). Not every method can process the 125,000 genomes included in LEMMI/RefSeq with the resources provided (245 GB of RAM, representing a medium-scale environment). It is therefore relevant both to assess which subsets constitute good tradeoffs in terms of runtime, memory use, and accuracy of the predictions, and to track the impact of continuous database growth on different methods^25^.

### In-silico datasets and genome exclusion

LEMMI uses its repository not only as source material provided to each method to create their reference database, but also for sampling mock microbial communities to generate in-silico paired-end short reads used to measure the accuracy of predictions made by each method. It implements a genome exclusion^26^ approach to simulate unknown organisms at various taxonomic ranks and to prevent overfitting by excluding the source of the analyzed reads from the reference (Supplementary Figure 4, see Online Methods). Simulating taxa that have no corresponding organisms in the reference also makes LEMMI datasets good proxies to real life scenarios and creates a challenging problem that can be useful to spot methods producing an excess of false positive matches (Figure 3C). Datasets generated in-house by LEMMI contain randomly sampled bacteria and archaea, representing generic communities of variable complexity in terms of number of species, abundance distribution (i.e. absence or presence of taxa below 100 reads), and k-mers content. Their description is presented in Supplementary Table 1. Their detailed taxonomic compositions remain private as long as they are in use in the platform and will be published when LEMMI moves to its next major release. A demo dataset is available to exemplify their typical composition and help preparing a method container for evaluation. LEMMI datasets are complemented by others used in previously published benchmarking studies (CAMI1^10^ and mockrobiota^27^). This enables comparisons with past results and introduces variety in the dataset creation methods, which is useful to identify behaviors inherent to the benchmarking design (Supplementary Note). Detailed evaluation results using up to 17 metrics are presented individually in interactive plots for each dataset and taxonomic rank, namely genus and species (Figure 3, Supplementary Figure 2-3, 5-9, Supplementary Table 2).

### Two benchmarking categories

To allow a separate interpretation of evaluations conducted with pre-processed databases (meaning no genome exclusion and thus potential overfits of some references) from those conducted under an identical set of genomes absent from the source of in-silico reads, LEMMI distinguishes two benchmarking categories (Figure 2). When choosing “TOOLS & REFERENCES”, users can get an overview of all tools and references that exist, giving an outline of the ability of currently available approaches to capture the known microbial diversity, strongly influenced by the database sampling strategy and creation date, but reflecting what practitioners will encounter when using ready-to-use solutions. The second category, “METHOD ALGORITHMS”, considers only methods and datasets that permit the creation of a reference, using genome exclusion, that will be identical for each run. This is the best possible benchmark for developers to support their algorithm improvement claims, and for advanced user interested in producing their own reference database after selecting the most efficient method.

### Evaluation of recent methods

The release of the LEMMI benchmarking platform described here (beta01.20191002) includes the well-established methods Centrifuge^1^, Kraken1^20^ and Kraken2^21^ associated with Bracken^22^, Kaiju^28^, and Metaphlan2^2^. They are a good representation of the field as it is today, as they stand among methods evaluated together recently^6^. This allows LEMMI users to judge additional methods never evaluated before side-by-side with tools they are already familiar with. We highlight on Figure 2, Figure 3, and Supplementary Figure 2, 5-8 a glimpse of the performances of the novel algorithms implemented by Ganon^29^, Metacache^30^, and CCMetagen^31^, which are described in their supporting publications but currently absent from comparative benchmarks. We report overall very good performances in recall and precision at the species level for Metacache and Ganon, which make them worth considering as alternatives to older solutions if their specific innovations, for instance the ability to update the database provided by Ganon, are beneficial to the user. These tools perform well when compared to other solutions using a freely selected references (Figure 2, Figure 3AB) as well as when compared to other tools using an identical built reference (Figure 3C, Supplementary Figure 5). They are also top scorers when taxonomic binning objectives are considered (Supplementary Figure 6). CCMetagen shows very good precision but suffers from a poor recall, except when analyzing the two datasets having a lower number of species (Figure 3ABC, Supplementary Figure 7). As LEMMI minor releases are continuously produced, the current list of methods and databases (Supplementary Table 3) will rapidly be enriched. To appreciate the full extent of the results through detailed plots, we invite readers to visit the LEMMI web platform (https://lemmi.ezlab.org).

### Evaluation of different references with Kraken2

Taking advantage of LEMMI flexible reference construction, we appreciated the improvement of the Kraken algorithm in its second version^21^ regarding its ability to build comprehensive references without using a considerable amount of memory. It is the most efficient to that aspect among the methods evaluated in LEMMI so far (Figure 3D, Supplementary Figure 8) and it remains among the top scoring methods when combined with Bracken 2.0, being the top scorer in relative abundance estimation (Figure 2-3, Supplementary Figure 2). This lower memory usage enables LEMMI to run Kraken2 using of all entries in LEMMI/RefSeq to be compared to using the pre-built MiniKraken database, a reference intended for low memory environments and created by subsampling k-mers on “Complete genomes” (CG) sequences only. The diversity of datasets offered by LEMMI shows that good performances of Minikraken depend almost entirely on the source of the in-silico reads being CG sequences. This way of limiting source genomes, unrelated to the Minikraken k-mers subsampling strategy, ignores the large part of the diversity proxied by NCBI species taxids that have no CG representatives (Supplementary Note). This can be observed when analyzing CAMI1 datasets, in which a large proportion of taxa is overlooked while it could actually be recovered using LEMMI/RefSeq as reference (Supplementary Figure 9).

## Discussion

Here, we introduce the first of its kind continuous integration benchmarking platform for metagenomics classifiers to enable immediate and independent evaluation of any newly published method using datasets of various compositions that can be generated or reused from previous effort. In addition, we highlight the high potential of some recently published methods unevaluated independently before, and ensure that future promising developments will join the ranking soon after they are identified as such. We also show the importance of considering all assembly states issued from RefSeq to avoid missing most of the diversity produced by year of microbial genome sequencing. Therefore, better subsampling strategies than simply keeping CG sequences should be explored by database creators to mitigate the problem of insufficient computational resources when conducting analyses. Moreover, future benchmarking efforts should not rely exclusively on CG to evaluate methods in order to assess the true ability of methods to scale with the ever-growing diversity of sequenced taxa.

LEMMI envisions a sustainable life cycle (Supplementary Figure 1b), encouraging feedback from the community of method developers and users. It ensures a traceable archive of past rankings through the use of unique “fingerprints” to be reported in publications. While the current major release of LEMMI uses the NCBI taxonomy^32^ and has put the focus on evaluating the lowest supported ranks, i.e. genus and species, it is designed to integrate alternative ranks or taxonomies. There are calls to revise NCBI taxonomy^33,34^ and future method implementations could be more agnostic regarding taxonomic authority (i.e. configuring the ranks and identifiers, instead of having the NCBI taxonomy entangled with the core classification algorithm). This would enable the exploration of new approaches meant to improve the classification resolution (e.g. reaching bacterial strains, viral operational taxonomic units^35^). New LEMMI datasets (e.g. also including long read technologies) will be produced for the next major release to replace those in use today, integrating newly discovered taxon or covering currently unsupported clades. Having all methods containerized will facilitate a systematic re-evaluation of all valuable ones. Developers interested in submitting their tools can visit https://lemmi.ezlab.org, where documentation, support, a discussion board, and evaluated containers are available. Future extensions of LEMMI may also include a standalone version to allow private assessment and help with development before submitting to the public platform for an independent evaluation.

Benchmarking has become a must-have requirement for publishing novel methods. To bring credibility and facilitate the adoption by their target audience, it is essential that methods appear side-by-side with established competitors in a trusted independent ranking. The technology of containerization has a strong future in the bioinformatics community. Therefore, LEMMI will encourage developers to consider biocontainers to disseminate their work and to standardize the results formats so users will obtain easy-to-use and stable implementations of up-to-date methods as they appear in the benchmark.

## Online Methods

### Release strategy

LEMMI rankings are identified by the major release version of the platform (e.g beta01) and the minor release date (e.g. 20191002). Until the next major release, the content of the LEMMI/RefSeq repository, the version of the NCBI taxonomy, and the datasets are frozen to a specific version that guarantees exact comparisons. Continuous additions or withdrawals of an entry in the rankings generate a new minor release date, while previous rankings remain accessible by requesting old fingerprints (e.g. https://lemmi.ezlab.org/#/rankings/SD.DEFAULT.beta01.20191002).

### Structure of the pipeline

An in-house python3-based^36,37^ controller coordinates the many subtasks required to generate datasets, run the candidate containers, and compute the statistics. Snakemake 5.3.1^38^ is used to supervise individual subtasks such as generating a dataset or running one evaluation. The process is semi-automated through configuration files, designed to allow a potential full automation through a web application. To be easily deployable, the benchmarking pipeline itself is wrapped in a Docker container. The plots presented on the user interface are generated with the mpld3 library^39^: https://mpld3.github.io.

### LEMMI containers

The LEMMI containers are implemented for Docker 18.09.0-ce. They partially follow the design^40^ introduced by http://bioboxes.org/ as part of the first CAMI challenge effort. The required output files are compatible with the profiling and binning format created for the CAMI challenge. Two tasks have to be implemented in order to generate a reference and conduct an analysis. To take part in the benchmark, a method developer has to build the container on their own environment, while ensuring that both tasks can be run by an unprivileged user and return the desired outputs. A tutorial is available on https://gitlab.com/ezlab/lemmi/wikis/user-guide. The containers or the sources to recreate them are made available to the users. The sources of all method containers presented as results in this study are available on https://gitlab.com/ezlab/lemmi/tree/beta01.20191002/containers.

### Computing resources

During the benchmarking process, the container is loaded on a dedicated server and given 245 GB of RAM and 32 cores. Reaching the memory limit will cause the container to be killed ending the benchmarking process unsuccessfully. All inputs and outputs are written on a local disk and the container is not given access to the Internet.

### Taxonomy

The NCBI taxonomy is used to validate all entries throughout the process and unknown taxids are ignored (unclassified). The framework etetoolkit^41^ (ETE3) is used to query the taxonomy. The database used in beta01 was downloaded on 03/09/2018 and remains frozen to this version until a new major release of the LEMMI platform.

### RefSeq repository

All RefSeq assemblies for bacteria, archaea, and viruses were downloaded from ftp.ncbi.nlm.nih.gov/refseq/release (download date for the beta01 release was 08/2018) with the conditions that they contained both a protein and a nucleotide file and that their taxid has a corresponding entry in the ETE3 NCBI taxonomy database, for a total of 132,167 files of each sequence type. The taxonomic lineage for the seven main levels was extracted with ETE3 (superkingdom, phylum, class, order, family, genus, species). Viruses were not used for generating references and datasets in this proof of concept study and the number of genomes in the reference RefSeq/08.2018/All is 124,289. To subset the repository and keep one representative per species as inputs for the reference construction (for entries labelled as RefSeq/08.2018/1rep.), the list of bacterial and archaeal genomes was sorted according to the assembly states (1:Complete Genome, 2:Chromosome, 3:Scaffold, 4:Contig) and the first entry for each species taxid was retained, for a total of 18,907 files. When subsampling the repository in the genome exclusion mode (i.e. for the METHOD ALGORITHMS category, Supplementary Figure 4), if the entry to be selected was part of the reads, the next representative in the list was used instead when available.

### LEMMI datasets

To sample the genomes included in the LEMMI datasets, a custom python script was used to randomly select representative genomes in the LEMMI/RefSeq assembly repository, among bacterial and archaeal content (Supplementary Table 1). Their abundance was randomly defined following a lognormal distribution (mean=1, standard deviation in Supplementary Table 1). In the case of LEMMI_LOWDIV datasets, additional low coverage species (abundance corresponding to < 100 reads) were manually defined while in LEMMI_MEDDIV datasets, low coverage species were produced as part of the random sampling procedure. Each species abundance was normalized according to the species average genome size (as available in the LEMMI/RefSeq repository) to get closer to organisms’ abundance (considering one genome copy per cell), and the total was normalized to one to constitute a relative abundance profile. Therefore, both tools classifying all reads and those using markers genes can normalize their output to provide a unified answer. BEAR^42^ was used to generate paired-end reads, 2×150 bp, and DRISEE^43^ was used to extract an error profile from the SRA entry ERX2528389 to be applied onto the generated reads. The ground truth profile for the seven taxonomic ranks, and taxonomic bins for species and genus were kept. The non-unique 50-mers and 31-mers diversity of the obtained reads were generated with Jellyfish 2.2.844 on the concatenated pair of reads using the following parameters: jellyfish count -m 31 -s 3G --bf-size 5G -t 8 -L 1 reads.fq.

### Additional datasets

The CAMI1 datasets were obtained from https://data.cami-challenge.org/ (accessed 09/2018) along with the metadata describing their content, already in the expected file format. The binning details were reprocessed to obtain distinct lists at the species and genus rank. The mockrobiota-17 dataset^45^ was obtained through https://github.com/caporaso-lab/mockrobiota and reprocessed to obtain a taxonomic profile in the appropriate format. No binning detail is available for this dataset, therefore no assessment of this aspect is based on this dataset. 50-mers and 31-mers diversity were computed as detailed above.

### Analysis of the results

The profile and binning reports are processed with OPAL 0.2.8^12^ and AMBER 0.7.0^13^ against the ground truth to obtain a wide range of metrics. Binning reports are processed to obtain a file for each taxonomic rank (genus/species), moving reads up from lowest level. The profiles of the candidate methods and the ground truth are filtered to discard low coverage taxa at different thresholds (below values corresponding to 1/10/100/1000 reads) and all metrics are computed for all values. When a container is not able to provide a profile as output, the LEMMI platform generates one using the proportion of reported reads. Taxa detection metrics are based on OPAL and thus on the profile output. Methods reporting a profile with 0.0 for low abundance taxa despite being present in their binning files will shift the balance from recall to precision. The low abundance score takes into account both the profile and binning output and is a custom metric calculated separately to evaluate the ability of the method to correctly identify organisms represented with very low read coverage, but penalizing methods likely to recover them by recurrent report of the same taxids owed to very poor precision. To achieve this, as precision of low abundance organisms cannot be defined for a single dataset (false positives always have a true abundance of zero and cannot be categorized as low abundance), the metric is computed by pairing two datasets to judge if a prediction can be trusted. The datasets (D1 and D2) include sets of taxa T1 and T2 that contain a subset of low abundance taxa (T1_low ≠ T2_low, < 100 reads coverage,). Each taxon belonging to T1_low identified in D1 increases the low abundance score given to the method for D1 (recall) only when it is not identified in D2 if absent from T2. Otherwise, a correct prediction of the taxon in D1 is canceled and does not improve the score (acting as proxy for low abundance precision). This is illustrated with Supplementary Figure 10. The score (0.0 - 1.0) is processed from both sides (D1, D2), to obtain an independent score for each of the paired dataset. This metric is only defined for the pair of LEMMI_LOWDIV and the pair of LEMMI_MEDDIV datasets (low abundance species: n=10, n=8, n=98, n=138 for LEMMI_LOWDIV_001, LEMMI_LOWDIV_002, LEMMI_MEDDIV_001, LEMMI_MEDDIV_002 respectively). The runtime corresponds to the time in minutes during which the container is loaded. The memory is the peak value of total_rss memory reported when the container is loaded to complete one task. Methods unable to deliver part of the expected outputs are assigned 0.0 for the corresponding metrics (e.g. methods unable to provide a read binning report).

### Ranking score

All metrics that are not already values between 0.0 and 1.0, with 1.0 being the best score, are transformed. The L1 distance is divided by its maximum value of 2.0 and subtracted from 1.0, the weighted UniFrac score is divided by its maximum value of 16.0 and subtracted from 1.0. The unweighted UniFrac score is divided by an arbitrary value of 25,000 and subtracted from 1.0. The memory and runtime are divided by 2x the maximum value (as defined by the LEMMI user through the interface) and subtracted from 1.0, to obtain a range between 0.5 and 1.0. This approach allows the user to segregate methods that remain below the limit from those that exceed it and get the value 0.0. Any transformed metric below 0.0 or above 1.0 is set to be 0 and 1 respectively. Each value is calculated for genus and species at 1, 10, and 100 reads low coverage filtering level and used or ignored according to the choice of the LEMMI user regarding these parameters. The final score displayed in the ranking is the harmonic mean of all metrics, taken into account 0, 1, or 3 times depending on the weight assigned to the metric by the LEMMI user.

## Data availability

All containers representing evaluated methods are available on https://lemmi.ezlab.org. The LEMMI demo dataset is available in the Zenodo repository with the identifier. https://zenodo.org/record/2651062.

## Acknowledgement

We would like to thank all members of the Zdobnov group for discussions and feedback. This work was supported by the Swiss National Science Foundation funding 31003A_166483 and 310030_189062 to E.Z.

## Author contributions

MS and EZ conceived the study. MS coded the platform. MS and MM conducted the analyses. MS and MM wrote the documentation. MS, MM and EZ wrote the manuscript.

## Competing interests

The authors declare no competing interests.

## Notes

#### Summary of Updates

The evaluation of recently published methods is discussed.

## Reference

1. Kim, D., Song, L., Breitwieser, F. P. & Salzberg, S. L. Centrifuge: rapid and sensitive classification of metagenomic sequences. Genome Res. 26, 1721–1729 (2016).

2. Truong, D. T. et al. MetaPhlAn2 for enhanced metagenomic taxonomic profiling. Nat. Methods 12, 902–903 (2015).

3. Jacobs, J. Microbe Land. (2018). https://microbe.land/2018/12/13/97-metagenomics-classifiers/

4. Norel, R., Rice, J. J. & Stolovitzky, G. The self-assessment trap: can we all be better than average? Mol. Syst. Biol. 7, 537 (2011).

5. McIntyre, A. B. R. et al. Comprehensive benchmarking and ensemble approaches for metagenomic classifiers. Genome Biol. 18, 182 (2017).

6. Ye, S. H., Siddle, K. J., Park, D. J. & Sabeti, P. C. Benchmarking Metagenomics Tools for Taxonomic Classification. Cell 178, 779–794 (2019).

7. Lindgreen, S., Adair, K. L. & Gardner, P. P. An evaluation of the accuracy and speed of metagenome analysis tools. Sci. Rep. 6, (2016).

8. Gardner, P. P. et al. Identifying accurate metagenome and amplicon software via a meta-analysis of sequence to taxonomy benchmarking studies. PeerJ 7, e6160 (2019).

9. Peabody, M. A., Van Rossum, T., Lo, R. & Brinkman, F. S. L. Evaluation of shotgun metagenomics sequence classification methods using in silico and in vitro simulated communities. BMC Bioinformatics 16, (2015).

10. Sczyrba, A. et al. Critical Assessment of Metagenome Interpretation—a benchmark of metagenomics software. Nat. Methods 14, 1063–1071 (2017).

11. Bremges, A. & McHardy, A. C. Critical Assessment of Metagenome Interpretation Enters the Second Round. mSystems 3, (2018).

12. Meyer, F. et al. Assessing taxonomic metagenome profilers with OPAL. Genome Biol. 20, (2019).

13. Meyer, F. et al. AMBER: Assessment of Metagenome BinnERs. GigaScience 7, (2018).

14. Belmann, P. et al. Bioboxes: standardised containers for interchangeable bioinformatics software. GigaScience 4, (2015).

15. Capella-Gutierrez, S. et al. Lessons Learned: Recommendations for Establishing Critical Periodic Scientific Benchmarking. (Bioinformatics, 2017). doi:10.1101/181677

16. Mangul, S. et al. Systematic benchmarking of omics computational tools. Nat. Commun. 10, 1393 (2019).

17. Maier-Hein, L. et al. Why rankings of biomedical image analysis competitions should be interpreted with care. Nat. Commun. 9, (2018).

18. Mangul, S., Martin, L. S., Eskin, E. & Blekhman, R. Improving the usability and archival stability of bioinformatics software. Genome Biol. 20, (2019).

19. da Veiga Leprevost, F. et al. BioContainers: an open-source and community-driven framework for software standardization. Bioinformatics 33, 2580–2582 (2017).

20. Wood, D. E. & Salzberg, S. L. Kraken: ultrafast metagenomic sequence classification using exact alignments. Genome Biol. 15, R46 (2014).

21. Wood, D. E., Lu, J. & Langmead, B. Improved metagenomic analysis with Kraken 2. (Bioinformatics, 2019). doi:10.1101/762302

22. Lu, J., Breitwieser, F. P., Thielen, P. & Salzberg, S. L. Bracken: estimating species abundance in metagenomics data. PeerJ Comput. Sci. 3, e104 (2017).

23. Truong, D. T. et al. MetaPhlAn2 for enhanced metagenomic taxonomic profiling. Nat. Methods 12, 902–903 (2015).

24. O’Leary, N. A. et al. Reference sequence (RefSeq) database at NCBI: current status, taxonomic expansion, and functional annotation. Nucleic Acids Res. 44, D733–D745 (2016).

25. Nasko, D. J., Koren, S., Phillippy, A. M. & Treangen, T. J. RefSeq database growth influences the accuracy of k-mer-based lowest common ancestor species identification. Genome Biol. 19, (2018).

26. Peabody, M. A., Van Rossum, T., Lo, R. & Brinkman, F. S. L. Evaluation of shotgun metagenomics sequence classification methods using in silico and in vitro simulated communities. BMC Bioinformatics 16, (2015).

27. Bokulich, N. A. et al. mockrobiota: a Public Resource for Microbiome Bioinformatics Benchmarking. mSystems 1, (2016).

28. Menzel, P., Ng, K. L. & Krogh, A. Fast and sensitive taxonomic classification for metagenomics with Kaiju. Nat. Commun. 7, (2016).

29. Piro, V. C., Dadi, T. H., Seiler, E., Reinert, K. & Renard, B. Y. ganon: continuously up-to-date with database growth for precise short read classification in metagenomics. bioRxiv 406017 (2018). doi:10.1101/406017

30. Müller, A., Hundt, C., Hildebrandt, A., Hankeln, T. & Schmidt, B. MetaCache: context-aware classification of metagenomic reads using minhashing. Bioinformatics 33, 3740–3748 (2017).

31. Marcelino, V. R. et al. CCMetagen: comprehensive and accurate identification of eukaryotes and prokaryotes in metagenomic data. bioRxiv 641332 (2019). doi:10.1101/641332

32. Federhen, S. The NCBI Taxonomy database. Nucleic Acids Res. 40, D136–D143 (2012).

33. Parks, D. H. et al. Selection of representative genomes for 24,706 bacterial and archaeal species clusters provide a complete genome-based taxonomy. (Microbiology, 2019). doi:10.1101/771964

34. Parks, D. H. et al. A standardized bacterial taxonomy based on genome phylogeny substantially revises the tree of life. Nat. Biotechnol. (2018). doi:10.1038/nbt.4229

35. Paez-Espino, D. et al. IMG/VR v.2.0: an integrated data management and analysis system for cultivated and environmental viral genomes. Nucleic Acids Res. 47, D678–D686 (2019).

36. McKinney, W. Data Structures for Statistical Computing in Python. 6 (2010).

37. Oliphant, T. E. Guide to NumPy. (2015).

38. Koster, J. & Rahmann, S. Snakemake--a scalable bioinformatics workflow engine. Bioinformatics 28, 2520–2522 (2012).

39. Hunter, J. D. Matplotlib: A 2D Graphics Environment. Comput. Sci. Eng. 9, 90–95 (2007).

40. Belmann, P. et al. Bioboxes: standardised containers for interchangeable bioinformatics software. GigaScience 4, (2015).

41. Huerta-Cepas, J., Serra, F. & Bork, P. ETE 3: Reconstruction, Analysis, and Visualization of Phylogenomic Data. Mol. Biol. Evol. 33, 1635–1638 (2016).

42. Johnson, S., Trost, B., Long, J. R., Pittet, V. & Kusalik, A. A better sequence-read simulator program for metagenomics. BMC Bioinformatics 15, S14 (2014).

43. Keegan, K. P. et al. A Platform-Independent Method for Detecting Errors in Metagenomic Sequencing Data: DRISEE. PLoS Comput. Biol. 8, e1002541 (2012).

44. Marçais, G. & Kingsford, C. A fast, lock-free approach for efficient parallel counting of occurrences of k-mers. Bioinformatics 27, 764–770 (2011).

45. Kozich, J. J., Westcott, S. L., Baxter, N. T., Highlander, S. K. & Schloss, P. D. Development of a Dual-Index Sequencing Strategy and Curation Pipeline for Analyzing Amplicon Sequence Data on the MiSeq Illumina Sequencing Platform. Appl. Environ. Microbiol. 79, 5112–5120 (2013).

