## Supplementary material for "LEMMI: A continuous benchmarking platform for metagenomics classifiers"

Evgeny M. Zdobnov[1]*

[1]Department of Genetic Medicine and Development, University of Geneva Medical School and Swiss Institute of Bioinformatics, Geneva, Switzerland

#### Supplementary Note, Figures, and Tables

### Supplementary Note

#### Assembly state is not an efficient criterion for subsampling RefSeq, for benchmarking or analysis purpose.

Assemblies deposited on the RefSeq repository are flagged with "Complete genome" (CG), "Chromosome", "Scaffold", or "Contig" to describe the completion state of sequencing and assembly procedures. It is likely that the accelerating production of new assemblies, enabled notably by metagenomics, leads to fewer new entries being polished enough to obtain the CG flag, decreasing the representativity of this category over time (41% of the species taxid in the LEMMI/RefSeq repository in mid-2018, only 19% when ignoring viruses, which are not used in the beta01 release). The first LEMMI datasets (LEMMI_LOWDIV and LEMMI_MEDDIV) were created using only CG sequences to work with the best representative sequences (Supplementary Table 1). We noticed that the Minikraken databases (obtained in October 2017, updated in November 2018 and April 2019 for Kraken 2) performed well on these, while being unable to recover most of the species in the CAMI1 datasets, in contrast to using Kraken 2 with all LEMMI/RefSeq genomes available in mid-2018, confirming that species found in CAMI1 datasets are represented in the RefSeq assembly repository (Supplementary Figure 9). We realized that regular versions of Kraken databases are built using only sequences with the CG state[1]. Eventually, LEMMI_HIGHDIV_201802_001 (Supplementary Table 2) was designed to account for the whole diversity found in RefSeq by sampling all assembly states. Overall, any reference based on CG, such as Minikraken, will obtain a medium score in LEMMI, performing very well on LEMMI_LOWDIV and LEMMI_MEDDIV and very poorly on CAMI1 datasets. Future LEMMI datasets will not permit such misrepresentation of the existing taxonomy.

### Supplementary Figures

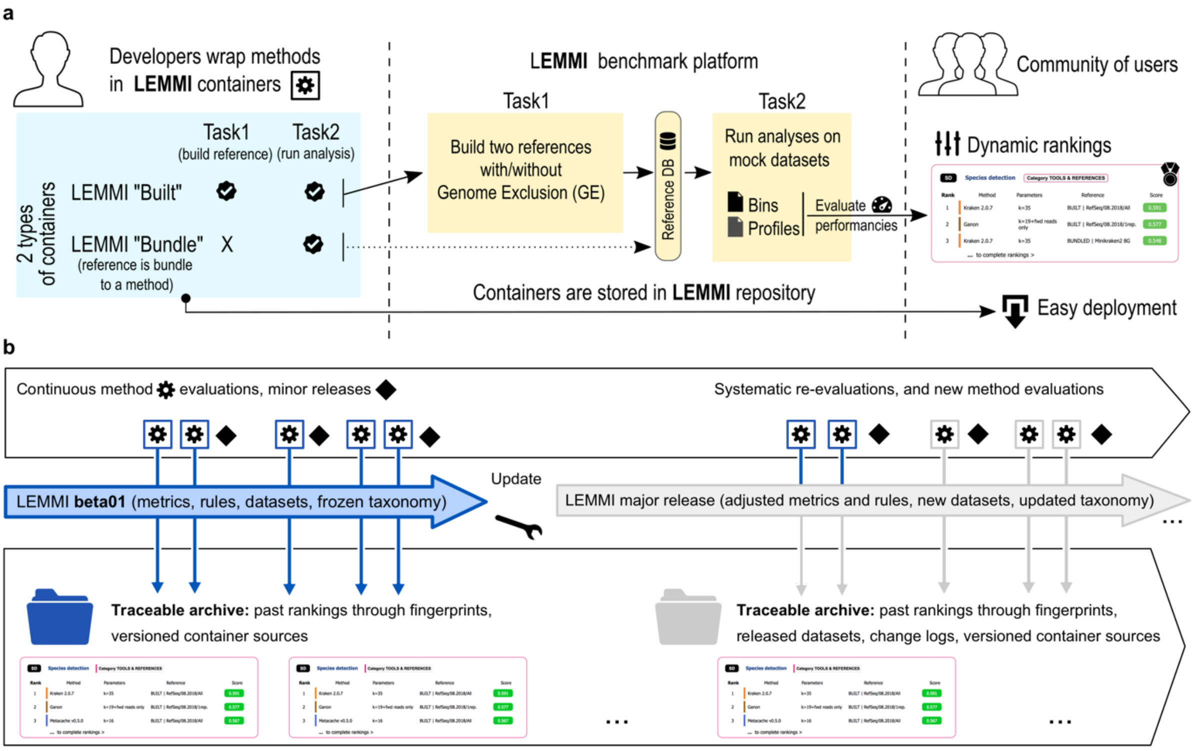


**Supplementary Fig. 1 | LEMMI workflow and expected life cycle.** (a) Developers prepare a container for their method following the provided guideline, to complete two tasks: building a reference using provided FASTA files (first task), and analyzing FASTQ samples to return a profile and binned reads (second task). Developers can also suggest a pre-packaged "bundled" reference instead of performing the first task. Their containerized method is then managed by the LEMMI administrator to be run within the LEMMI platform to process all datasets required to appear in the ranking. Multiple runs to explore parameters and references can be conducted using a single container. Method users can browse the results to define which methods best suit their needs and obtain the corresponding containers to conduct their own tests or actual analyses, with the guarantee of unified file formats and similar behaviors. (b) The release beta01 of LEMMI is the first major release. Every successful evaluation is integrated into the rankings, which are traceable through time and for which the source of the container is publicly available. Feedback from the community and progress in the field will eventually lead to the end of this first release to allow an update of both the platform and the datasets. While entering the next major release, still relevant methods will be systematically re-evaluated and new submissions will continue to populate the rankings.

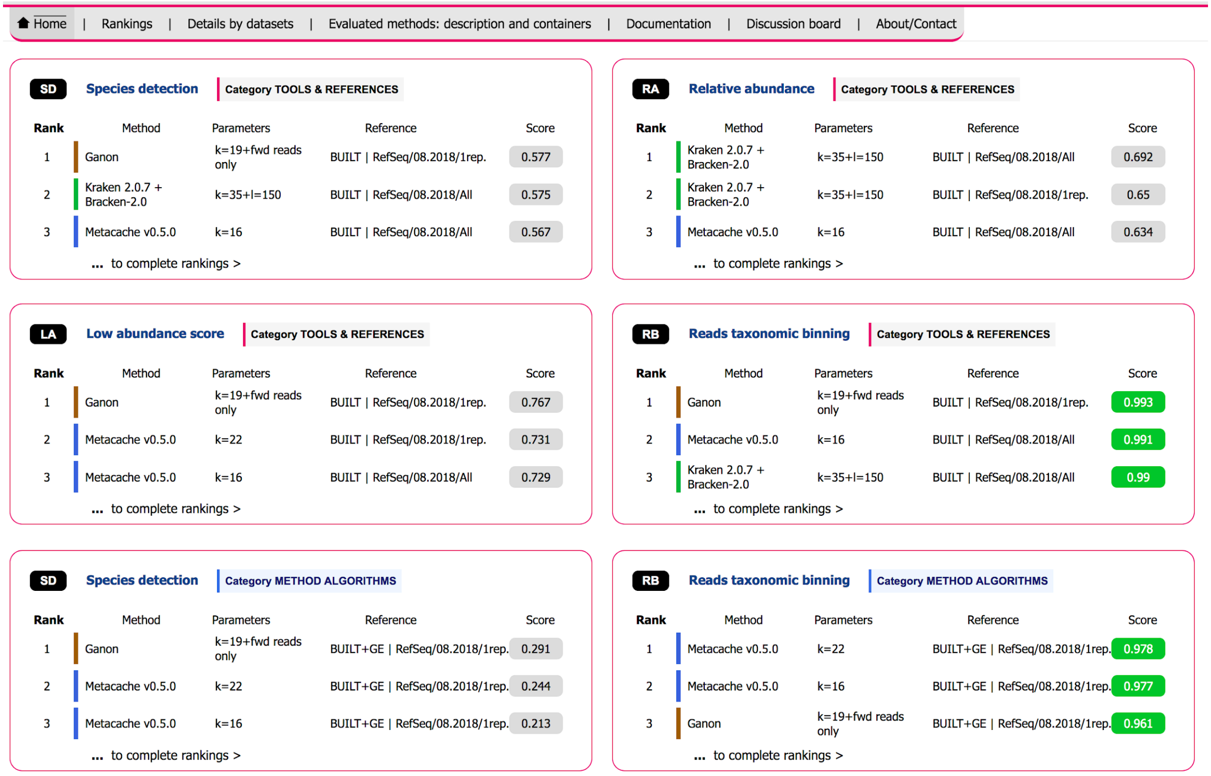


**Supplementary Fig. 2 | The LEMMI homepage.** The main page presents multiple rankings, each corresponding to the top three configurations (methods associated with a reference and specific parameters) according to metrics chosen for addressing various experimental objectives in one of the benchmark categories. These lists constitute different entry points to the dynamic ranking page where the full repertoire of configurations can be explored beyond these predefined criteria.

a.

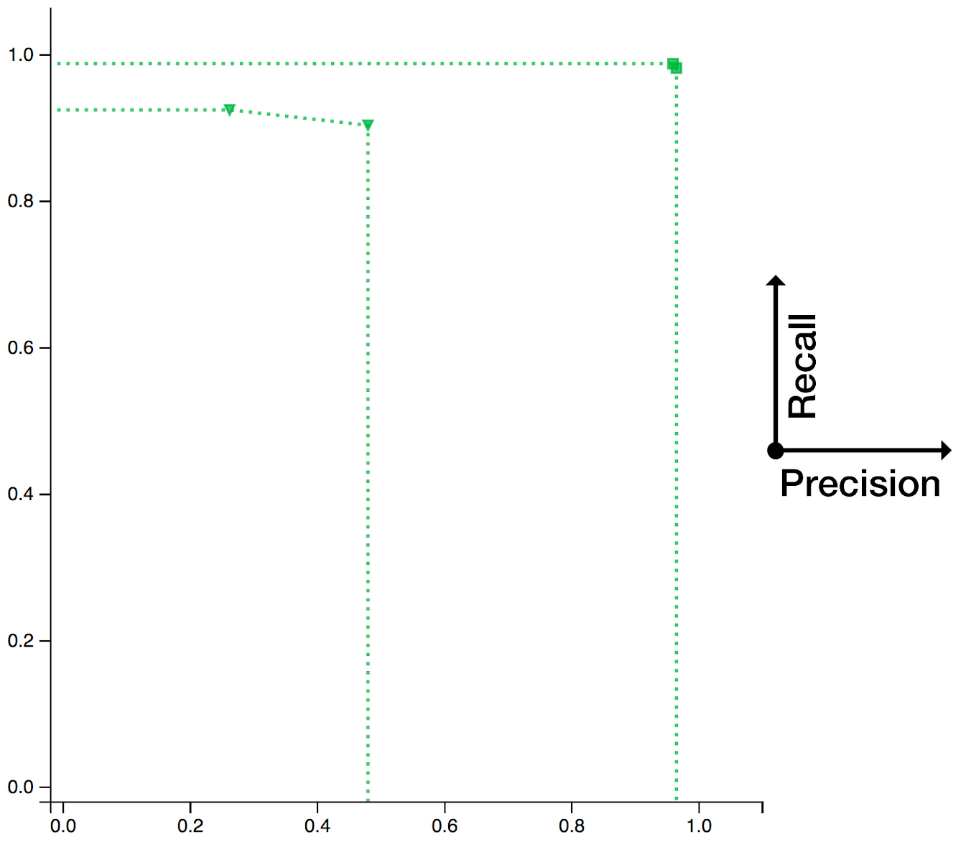

1.0
0.8
0.6
0.4
0.2
0.0
0.0
0.2
0.4
0.6
0.8
1.0
Recall
Precision


b.

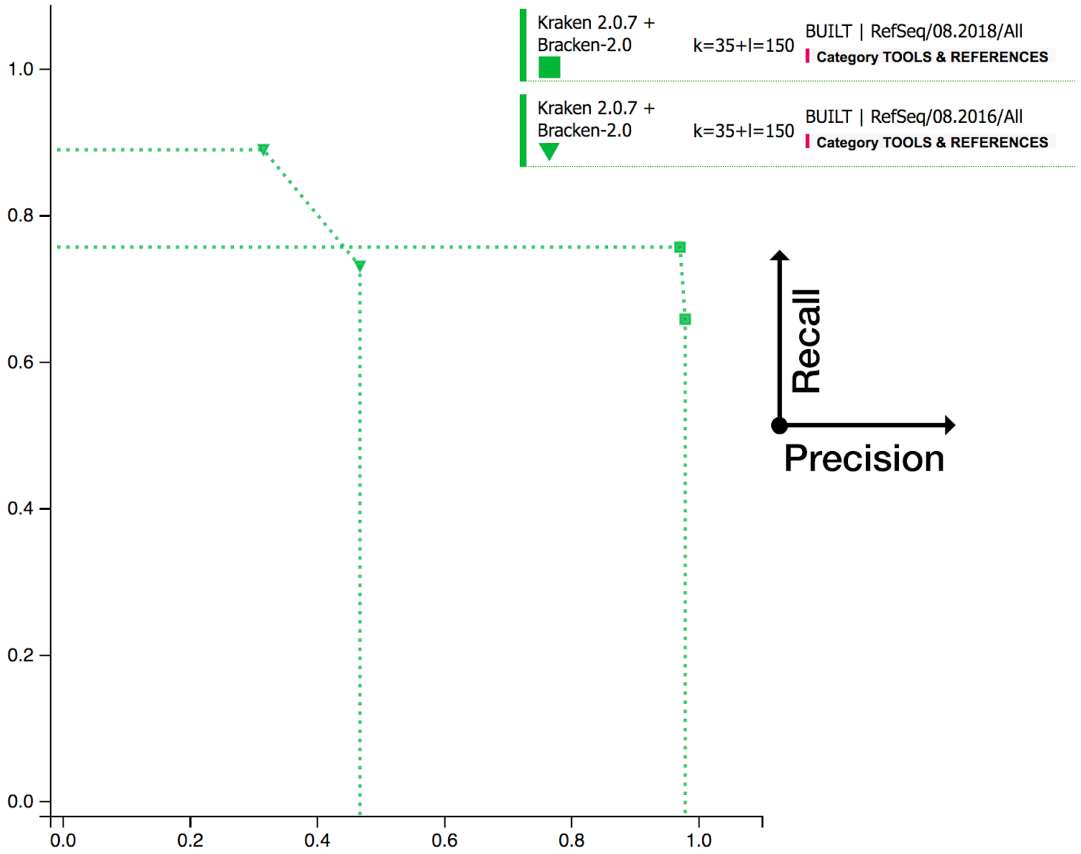

Kraken 2.0.7 + Bracken-2.0
k=35+l=150
BUILT | RefSeq/08.2018/All
Category TOOLS & REFERENCES
Kraken 2.0.7 + Bracken-2.0
k=35+l=150
BUILT | RefSeq/08.2016/All
Category TOOLS & REFERENCES
1.0
0.8
0.6
0.4
0.2
0.0
0.0
0.2
0.4
0.6
0.8
1.0
Recall
Precision

**Supplementary Fig. 3 | Using LEMMI as a reference time machine.** Kraken 2 analyses using the complete archaeal and bacterial content of the LEMMI repository from mid-2018, versus its state two years earlier. These two years doubled the number of genomes available as references. The taxonomic rank presented here is genus. An increase in the database size is not always beneficial in terms of recall, as reported previously[2], but is beneficial in terms of precision. Overall, selecting the most up to date reference remains necessary to cover newly sampled taxa, as reflected by the ranking on main Figure 2. (a) Precision-recall curve in species identification for the dataset LEMMI MEDIUM 1 (b) Precision-recall curve in species identification for the dataset LEMMI HIGH 1.

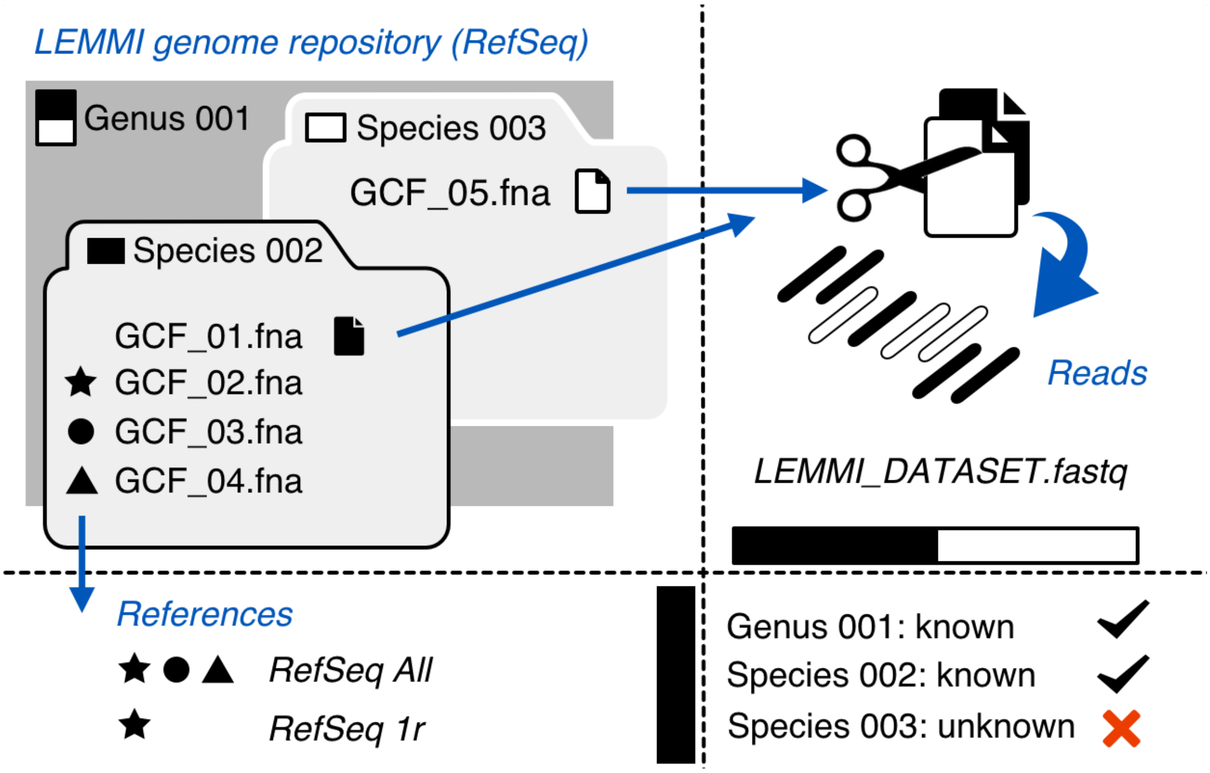


**Supplementary Fig. 4 | Creating unknown taxa from public data.** A toy example illustrating the "genome exclusion" approach used on LEMMI in-house datasets to avoid overfitting (i.e. having the source of the reads in the reference). Species 002 (black) and 003 (white) belonging to genus 001 have four representatives and one representative, respectively. One genome is taken out from each species to simulate the reads, and the rest is used to build a comprehensive reference (RefSeq All) or a reduced one (RefSeq 1r) using one randomly selected representative per species. All candidate methods provided with this scenario are expected to identify the genus 001 and the species 002. Species 003 becomes an unknown species, with no sequence available as reference.

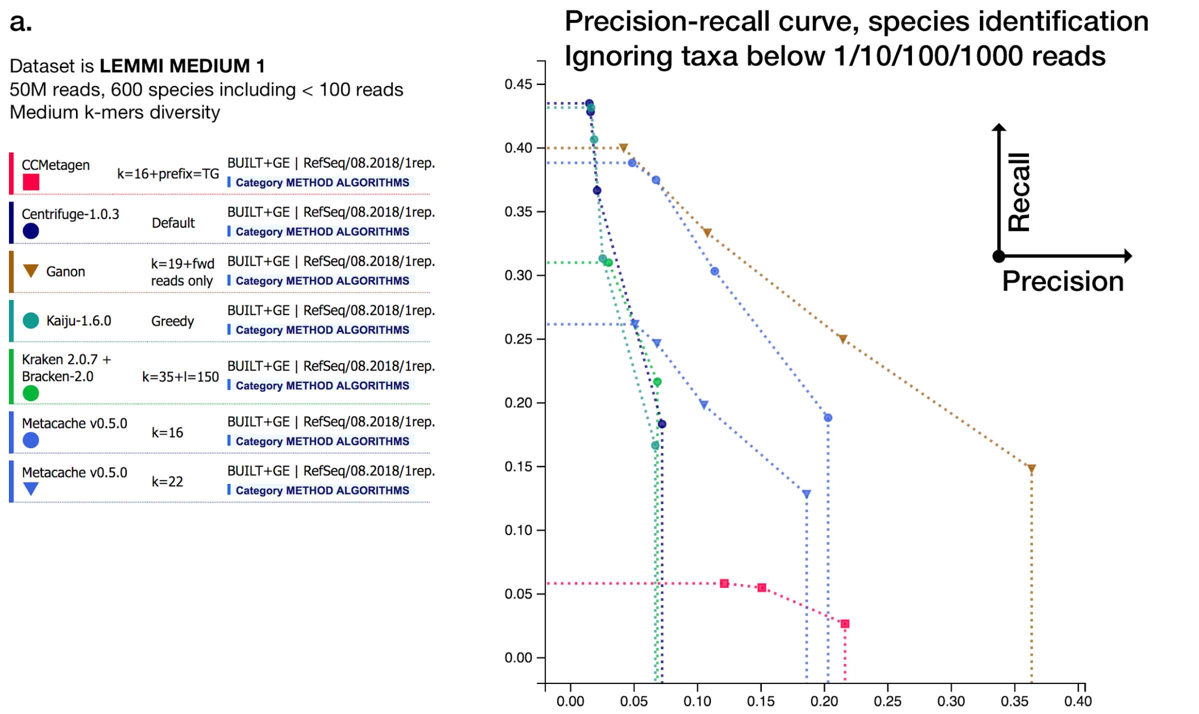


**b.**

| Rank | Method | Parameters | Reference ( ? ) | Score ( ? ) |
|---|---|---|---|---|
| 1 | Ganon | k=19+fwd reads only | BUILT+GE \| RefSeq/08.2018/1rep. | 0.291 |
| 2 | Metacache v0.5.0 | k=22 | BUILT+GE \| RefSeq/08.2018/1rep. | 0.244 |
| 3 | Metacache v0.5.0 | k=16 | BUILT+GE \| RefSeq/08.2018/1rep. | 0.213 |
| 4 | Centrifuge-1.0.3 | Default | BUILT+GE \| RefSeq/08.2018/1rep. | 0.113 |
| 5 | Kraken 2.0.7 + Bracken-2.0 | k=35+l=150 | BUILT+GE \| RefSeq/08.2018/1rep. | 0.106 |
| 6 | Kaiju-1.6.0 | Greedy | BUILT+GE \| RefSeq/08.2018/1rep. | 0.099 |
| 7 | CCMetagen | k=16+prefix=TG | BUILT+GE \| RefSeq/08.2018/1rep. | 0.071 |

**Supplementary Fig. 5 | Evaluation using an identical reference.** (a) Precision-recall curve in species identification of a mix of method using identical references built using one representative genome per species taxid, excluding the source of the reads. Not all species can be recovered as their only representative was used to produce reads. Therefore, the best recalls illustrated here are close to the maximum that can be reached at the species level under this scenario. (b) Ranking of methods over all datasets under the scenario described above given the LEMMI preset “Species detection”. Precision and recall are considered equally and taxa represented by less than 100 reads are ignored.

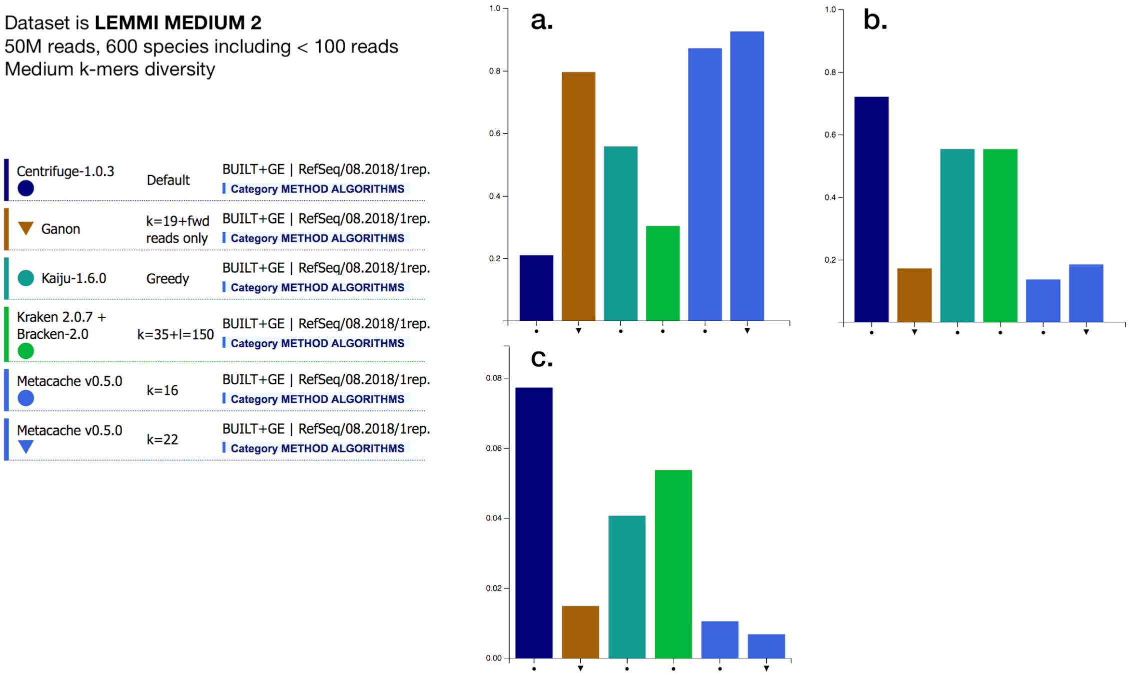


**Supplementary Fig. 6 | Taxonomic binning.** (a) Normalized rand index in species binning for a mix of methods using identical references built using one representative genome per species taxid, excluding the source of the reads. (b) Proportion of classified reads at the species level. (c) Proportion of reads assigned to a false positive species.

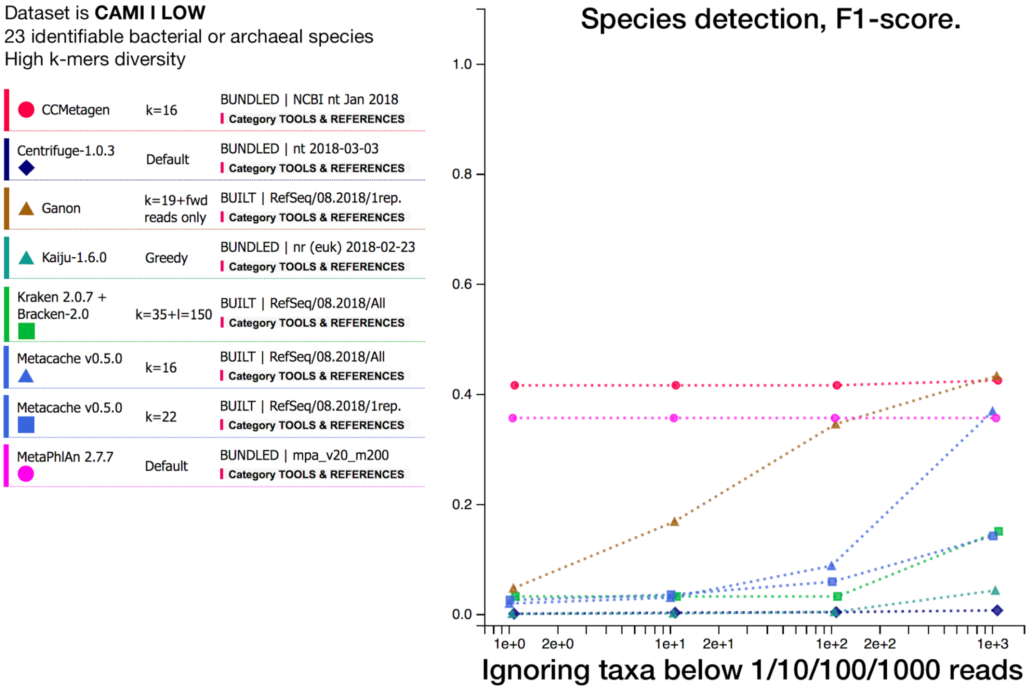


**Supplementary Fig. 7 | A dataset with a low number of species.** F1-score in species identification of a mix of method using freely provided references or built using the maximal capacity of the tool with 245GB. The dataset is the low complexity set from the first CAMI challenge.

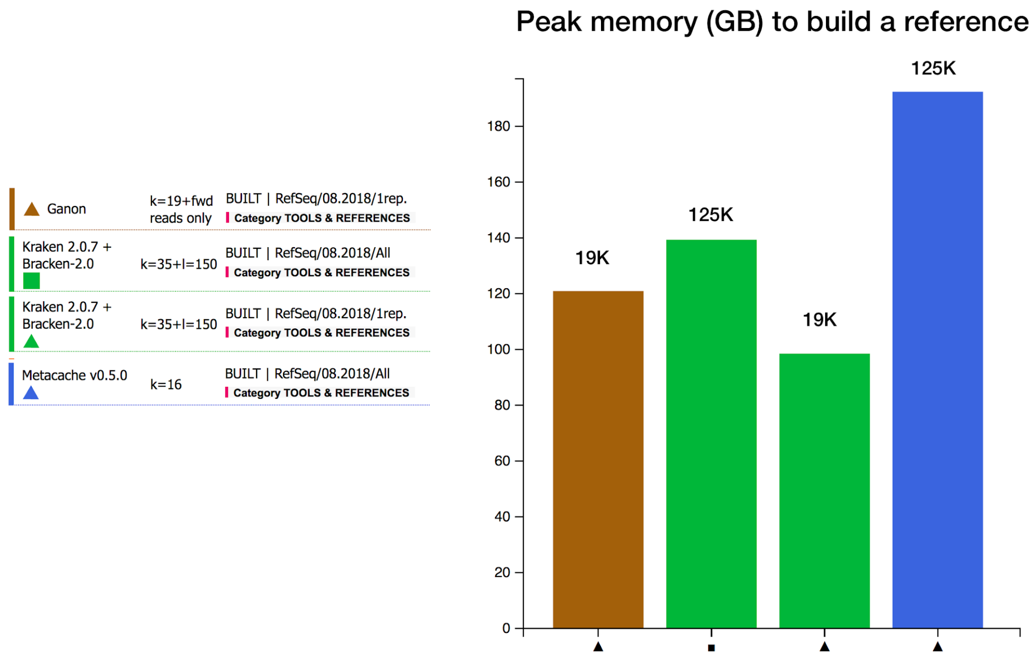


**Supplementary Fig. 8 | Resources usage.** Amount of memory used by Ganon, Kraken2, and Metacache to construct their reference. The number of genomes included, one representative per species (~19,000 files) or all representatives (~125,000 files) is indicated.

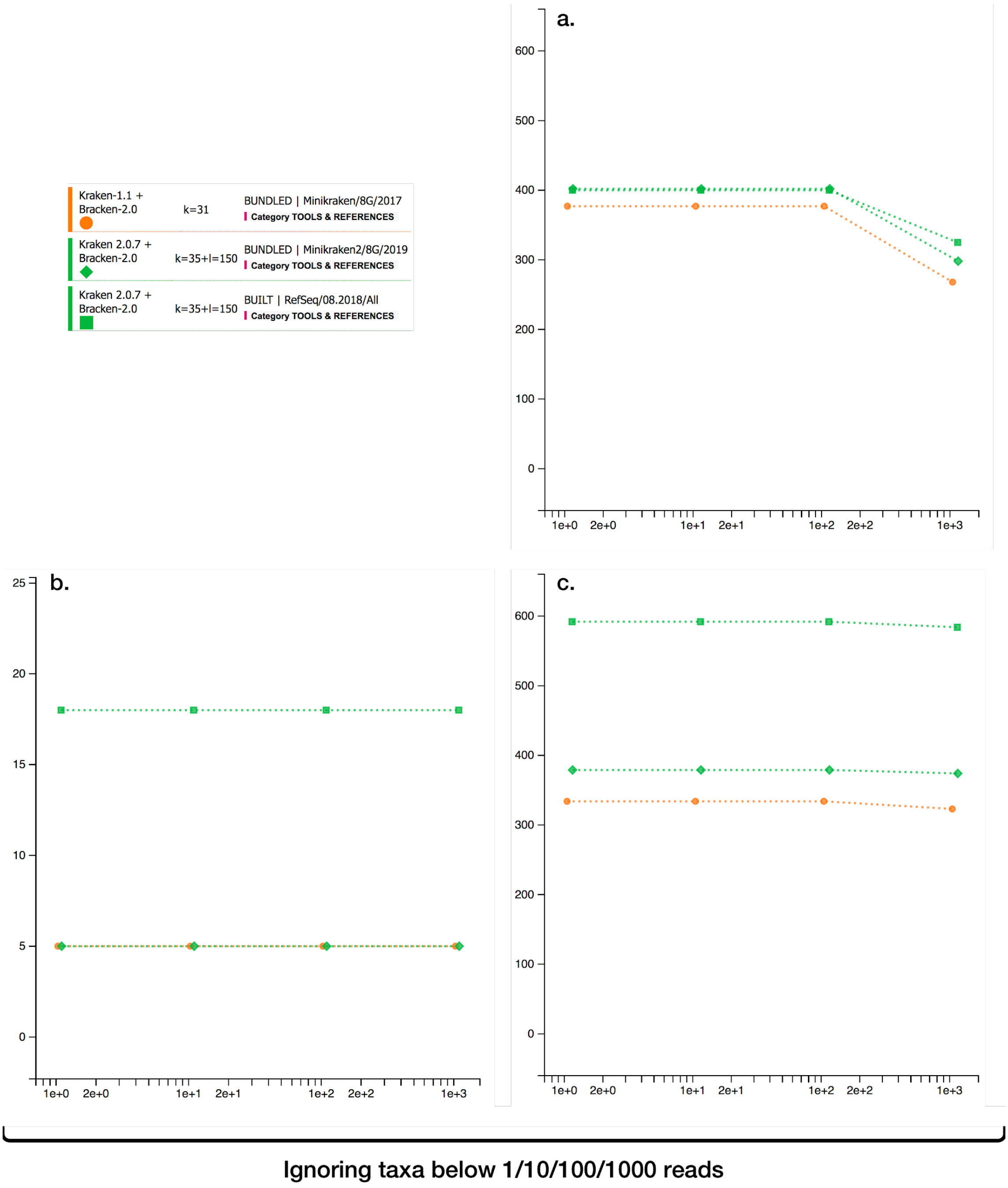


**Supplementary Fig. 9 | How the evaluation of the Minikraken database is biased by the design of the datasets used.** Counts of species correctly identified when running kraken1 and kraken2 with either Minikraken 2017/2019 or a comprehensive built of the LEMMI/RefSeq repository (mid-2018) without excluding any genome. (a) The dataset is LEMMI MEDIUM 1, based only on assemblies flagged as “Complete Genome”. (b) The dataset is CAMI1 LOW. (c) The dataset is LEMMI HIGH 1, sampled using all assembly states in the LEMMI/RefSeq repository. The latter can be seen as the fairest evaluation of the Minikraken database.

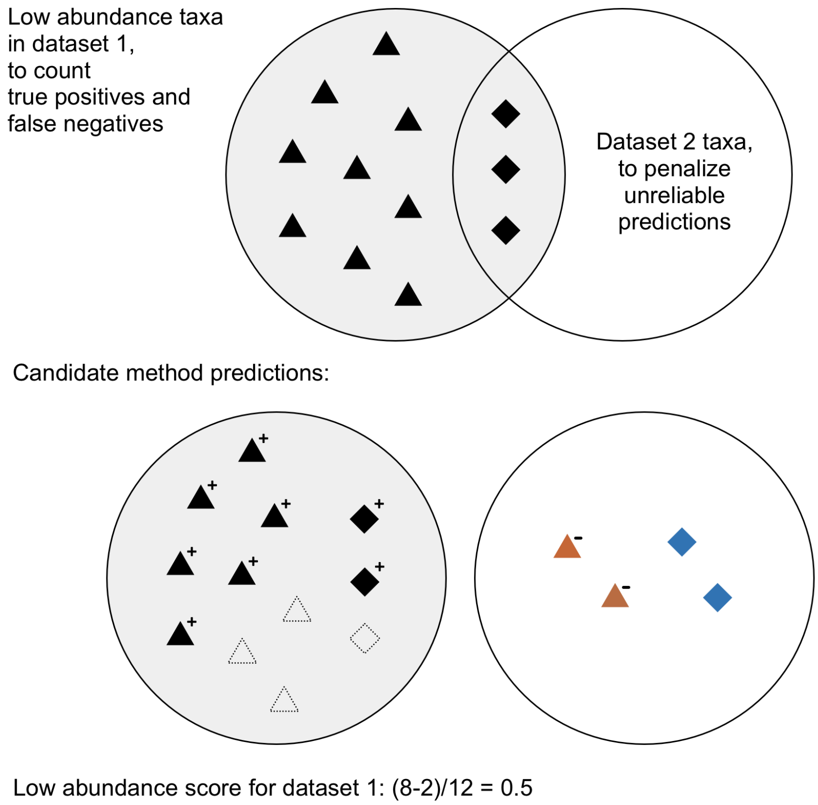


**Supplementary Fig. 10 | Low abundance score from the perspective of one of the paired datasets.** Triangles stand for species unique to the dataset 1 that is being evaluated and should not be found in the paired dataset 2. Diamonds represent species that are present in both datasets and will not affect the precision if found in dataset 2.

### Supplementary Tables

**Supplementary Table 1 |** Features of the datasets included in the LEMMI initial release. In the case of the LEMMI sets, unknown species and genera represent taxa for which all representatives were selected to generate the reads. Therefore, they are excluded from the reference when using genome exclusion (i.e. for the METHOD ALGORITHMS category). In other datasets, unknown taxa are those not found in the LEMMI/RefSeq repository dated from mid-2018. Datasets having taxa with less than 100 reads are used to compute the low abundance score.

| Name | Species count | Genera count | Unknown species | Unknown genera | < 100 reads species | < 100 reads genera | Non unique 50-mers | Number of reads | Abundance standard deviation | RefSeq assembly states |
|---|---|---|---|---|---|---|---|---|---|---|
| CAMI_I_LOW | 23 | 22 | 5 | 1 | 0 | 0 | 579,605,460 | 50M | - | - |
| CAMI_I_HIGH_1 | 243 | 194 | 86 | 7 | 0 | 0 | 1,496,568,850 | 50M | - | - |
| mockrobiota-17 | 10 | 18 | 0 | 0 | 0 | 0 | 68,345,393 | 1.2M | - | - |
| LEMMI_LOWDIV _201805_001 | 100 | 72 | 0 | 0 | 10 | 9 | 115,211,506 | 10M | 2.75 | Complete genome |
| LEMMI_LOWDIV _201805_002 | 100 | 71 | 0 | 0 | 8 | 4 | 105,634,790 | 10M | 2.75 | Complete genome |
| LEMMI_MEDDIV _201902_001 | 600 | 346 | 338 | 30 | 138 | 53 | 383,971,742 | 50M | 3.0 | Complete genome |
| LEMMI_MEDDIV _201902_002 | 600 | 339 | 332 | 40 | 98 | 44 | 455,319,422 | 50M | 3.0 | Complete genome |
| LEMMI_HIGHDIV _201902_001 | 600 | 333 | 393 | 34 | 2 | 2 | 1,340,621,833 | 50M | 1.75 | All |

**Supplementary Table 2 |** List of metrics available in the dataset detail pages (some datasets do not produce all of them). Underlined entries correspond to those contributing to the rankings.

| Metric | Comment |
|---|---|
| Taxa detection, precision-and-recall curve | Four data points corresponding to filtering abundance below 1/10/100/1000 reads<br>It comes with a second plot showing the area under the curve. |
| Taxa detection, recall | Four data points corresponding to filtering abundance below 1/10/100/1000 reads |
| Taxa detection, precision | Four data points corresponding to filtering abundance below 1/10/100/1000 reads |
| Taxa detection, true positive count | Four data points corresponding to filtering abundance below 1/10/100/1000 reads |
| Taxa detection, false positive count | Four data points corresponding to filtering abundance below 1/10/100/1000 reads |
| Taxa detection, F1-Score | Four data points corresponding to filtering abundance below 1/10/100/1000 reads |
| Unweighted UniFrac | The lower the better. Not specific to a taxonomic rank |
| Proportion of assigned reads | At the evaluated taxonomic rank or lower. No distinction of correct or incorrect assignment here. |
| Normalized rand index | Clustering accuracy at the evaluated taxonomic rank. Lower assignments are moved up to the evaluated rank for evaluation. |
| Proportion of reads assigned to a false positive taxa | At the evaluated taxonomic rank. Lower assignments are moved up to the evaluated rank for evaluation. Wrong assignment among true positive taxa are not included |
| Relative abundance error: L1 distance | |
| Weighted UniFrac | The lower the better. Not specific to a taxonomic rank |
| Low abundance score | Only for pair of LEMMI datasets. See methods and Supplementary Figure 10 |
| Runtime for analysis | For everything that the task within the container need to do, including cleaning/preprocessing fastq |
| Runtime for building the reference | For everything that the task within the container need to do, including cleaning/preprocessing fasta |
| Memory used for analysis | Peak memory in GB |
| Memory used for building the reference | Peak memory in GB |

**Supplementary Table 3 |** List of configurations (methods associated with a reference and specific parameters) included in release beta01.20191002 of the LEMMI platform. The configurations that successfully built a reference based on the entire LEMMI/RefSeq repository when provided with 245 GB of RAM are underlined. All others were limited to using one representative per species taxid. The source of corresponding containers can be found on https://gitlab.com/ezlab/lemmi/tree/beta01.20191002/containers

| **Method** | **Parameters** | **Reference** | **Benchmark category** | **Note** |
|---|---|---|---|---|
| Kaiju 1.6.0 | default | nr_euk 2018-02-23, bundled | TOOLS & REF. | |
| Centrifuge 1.0.3 | default | nt 2018-03-03, bundled | TOOLS & REF. | |
| Kraken 1.1<br>+ Bracken 2.0 | default | Minikraken 8G 2017, bundled | TOOLS & REF. | 125-mers db for bracken |
| Kraken 2.0.7<br>+ Bracken 2.0 | k=35 | Minikraken 8G 2019, bundled | TOOLS & REF. | 150-mers db for bracken |
| Kraken 2.0.7<br>+ Bracken 2.0 | k=35 | RefSeq/08.2018/All, built | TOOLS & REF. | 150-mers db for bracken |
| Kraken 2.0.7<br>+ Bracken 2.0 | k=35 | RefSeq/08.2018/1rep., built | TOOLS & REF. | 150-mers db for bracken |
| Kraken 2.0.7<br>+ Bracken 2.0 | k=35 | RefSeq/08.2016/All, built | TOOLS & REF. | 150-mers db for bracken |
| Metacache 0.5.0 | k=16 | RefSeq/08.2018/All, built | TOOLS & REF. | |
| Metacache 0.5.0 | k=22 | RefSeq/08.2018/1rep., built | TOOLS & REF. | |
| MetaPhlAn 2.7.7 | default | Mpa_v20_m200, bundled | TOOLS & REF. | |
| Ganon | k=19 | RefSeq/08.2018/1rep., built | TOOLS & REF. | Using only the forward reads |
| CCMetagen | k=16<br>+prefix TG | NCBI nt Jan 2018, bundled | TOOLS & REF. | |
| Kaiju 1.6.0 | default | RefSeq/08.2018/1rep.<br>Built with genome exclusion | METHOD ALGO. | |
| Centrifuge 1.0.3 | default | RefSeq/08.2018/1rep.<br>Built with genome exclusion | METHOD ALGO. | Limited to 12 cpus to build the reference |
| Kraken 2.0.7<br>+ Bracken 2.0 | k=35 | RefSeq/08.2018/1rep.<br>Built with genome exclusion | METHOD ALGO. | 150-mers db for bracken |
| Metacache 0.5.0 | k=16 | RefSeq/08.2018/1rep.<br>Built with genome exclusion | METHOD ALGO. | |
| Metacache 0.5.0 | k=22 | RefSeq/08.2018/1rep.<br>Built with genome exclusion | METHOD ALGO. | |
| Ganon | k=19 | RefSeq/08.2018/1rep.<br>Built with genome exclusion | METHOD ALGO. | Using only the forward reads |
| CCMetagen | k=16<br>+prefix TG | RefSeq/08.2018/1rep.<br>Built with genome exclusion | METHOD ALGO. | |

### References

1. Wood, D. E. & Salzberg, S. L. Kraken: ultrafast metagenomic sequence classification using exact alignments. *Genome Biol.* **15**, R46 (2014).
2. Nasko, D. J., Koren, S., Phillippy, A. M. & Treangen, T. J. RefSeq database growth influences the accuracy of k-mer-based lowest common ancestor species identification. *Genome Biol.* **19**, (2018).
